# Critical transitions and evolutionary hysteresis in movement: Habitat fragmentation can cause abrupt shifts in dispersal that are difficult to revert

**DOI:** 10.1101/2022.10.10.510956

**Authors:** Monique de Jager, Merel Soons

## Abstract

Under habitat fragmentation, plant species’ survival hinges on the ability of individuals to disperse from one habitat patch to another. While there is evidence that severe habitat fragmentation leads to evolution of reduced dispersal ability and that such decreased mobility is generally detrimental for species’ survival, it is unknown whether species adapt via a gradual loss in dispersal ability or via a sudden shift from frequent to infrequent dispersal between patches (i.e., a critical transition). Using a spatially explicit, individual-based modelling approach, we show that a small increase in inter-patch distance can generate an abrupt shift in plant seed dispersal strategy from long to short distances. Most importantly, we found that a substantial increase in connectivity between habitat fragments is required to reverse this loss of long-distance dispersal, due to an evolutionary hysteresis effect. Our theory prompts for re-consideration of the eco-evolutionary consequences of habitat fragmentation as restoring habitat connectivity may require restoration of much higher connectivity levels than currently assumed.

## Introduction

Globally, habitat fragmentation is a major cause of population declines and biodiversity loss (Diffenbaugh and Field 2013; IPBES 2019). Currently, 37% of grasslands, 70% of forests and 60% of the largest rivers are fragmented (Revenga et al. n.d.; Keeler-Wolf 2001; Haddad et al. 2015), with remaining natural habitat patches becoming smaller, fewer and farther apart, resulting in unprecedented rates of species extinction (Tilman et al. 2017; IPBES 2019). In this increasingly fragmented world, nature conservation and restoration activities often concentrate on reinstating connections between patches by securing and restoring nearby suitable locations or implementing corridors. Yet, such efforts are often ineffective (Race and Fonseca 1996; Suding 2011). When habitat conditions are sufficiently restored, the lack of success in restoring species populations is often attributed to insufficient nearby source populations to initiate recolonization (Donath et al. 2003; Brederveld et al. 2011; Sundermann et al. 2011). However, when quantified, the availability of source populations in the surrounding landscape only explains the lack of recolonization to a limited extent (e.g., ca. 20% in restored streams (Brederveld et al. 2011)) and other, unknown mechanisms are likely to also contribute substantially. It is of critical importance to identify these mechanisms, as failure of species to recolonize translates into failure to restore biodiversity and biodiversity-dependent ecosystem services (Isbell et al. 2015).

Restoration activities are based on the assumption that species’ dispersal abilities have been unaffected by the fragmented situation, leaving remaining populations well equipped to (re)colonize restored habitat patches (Honnay et al. 2002). Yet, dispersal is fundamental to an organisms fitness and is therefore under strong selection (Baguette et al. 2013). While habitat fragmentation can affect organisms in many ways, accumulating theoretical and empirical evidence points out that increased habitat fragmentation may lead to the evolution of short-distance dispersal (Cody and Overton 1996; Gandon and Michalakis 1999; Travis and Dytham 1999; Heino and Hanski 2001; Cheptou et al. 2008, 2017; Travis et al. 2010; Lindenmayer and Fischer 2013; Legrand et al. 2017). In highly fragmented conditions, (rapid) evolution of short-distance dispersal strategies increases dispersal within the parental patch while reducing the proportion of propagules lost to outside, uninhabitable areas. However, due to such a loss of dispersal capacity, colonization of new areas becomes impossible. Populations may thus become extremely vulnerable to local disturbances, gene flow between populations impaired, and networks of interactions with other species weakened (Leibold et al. 2004; Ronce 2007; Bonte and Dahirel 2017). In order to forecast this kind of eco-evolutionary dynamics and mitigate its effects through management and planning (Travis et al. 2010; Tilman et al. 2017), crucial knowledge is required regarding how altering habitat fragmentation affects the evolution of dispersal strategies.

In plants, an increasing body of evidence suggests that populations in long-term fragmented landscapes have indeed lost the ability to disperse over long distances (Cheptou et al. 2008, 2017; Travis et al. 2010; Lindenmayer and Fischer 2013; Legrand et al. 2017). One reason for this may be the loss or modification of dispersal vectors in fragmented landscapes (Cordeiro and Howe 2001; Ozinga et al. 2009). Another reason may be genetic loss of dispersal ability (Ronce 2007; Cheptou et al. 2008), either resulting from inbreeding depression or genetic drift due to small population sizes (Cheptou and Donohue 2011) or caused by evolutionary modification of seed dispersal traits in plant populations inhabiting fragmented landscapes (Cheptou et al. 2008, 2017).

We theorize that if loss of dispersal has a genetic basis, ongoing habitat fragmentation and restoration can lead to a hysteresis effect in dispersal evolution (Fig. 1). Increasing fragmentation may result in a tipping point in dispersal capacity: once long-distance dispersers can no longer colonize other habitat patches, those organisms that disperse over short distances and remain within the boundaries of the source population are more likely to survive, given the fact that far-dispersed propagules are lost to uninhabitable areas (Balkau and Feldman 1973; Hastings 1983; Holt 1985; Gandon and Michalakis 1999). Within a community, this process leads to short-distance dispersing species outcompeting long-distance dispersing species (Liao et al. 2013*b*, 2013*a*), and within a population, mutations resulting in short-distance dispersal suddenly become advantageous (Cheptou et al. 2008). We hypothesize that a long-distance dispersal strategy can abruptly evolve into short-distance (within-patch dispersal), once habitat patches become unreachable. Once local dispersal has evolved, we hypothesize that a substantial decrease in the distance between habitable areas is required to reverse the loss of dispersal capacity (a hysteresis effect; Fig. 1), as gradual changes in dispersal capacity may not evolve when habitable patches remain out of reach.

**Figure 1:**
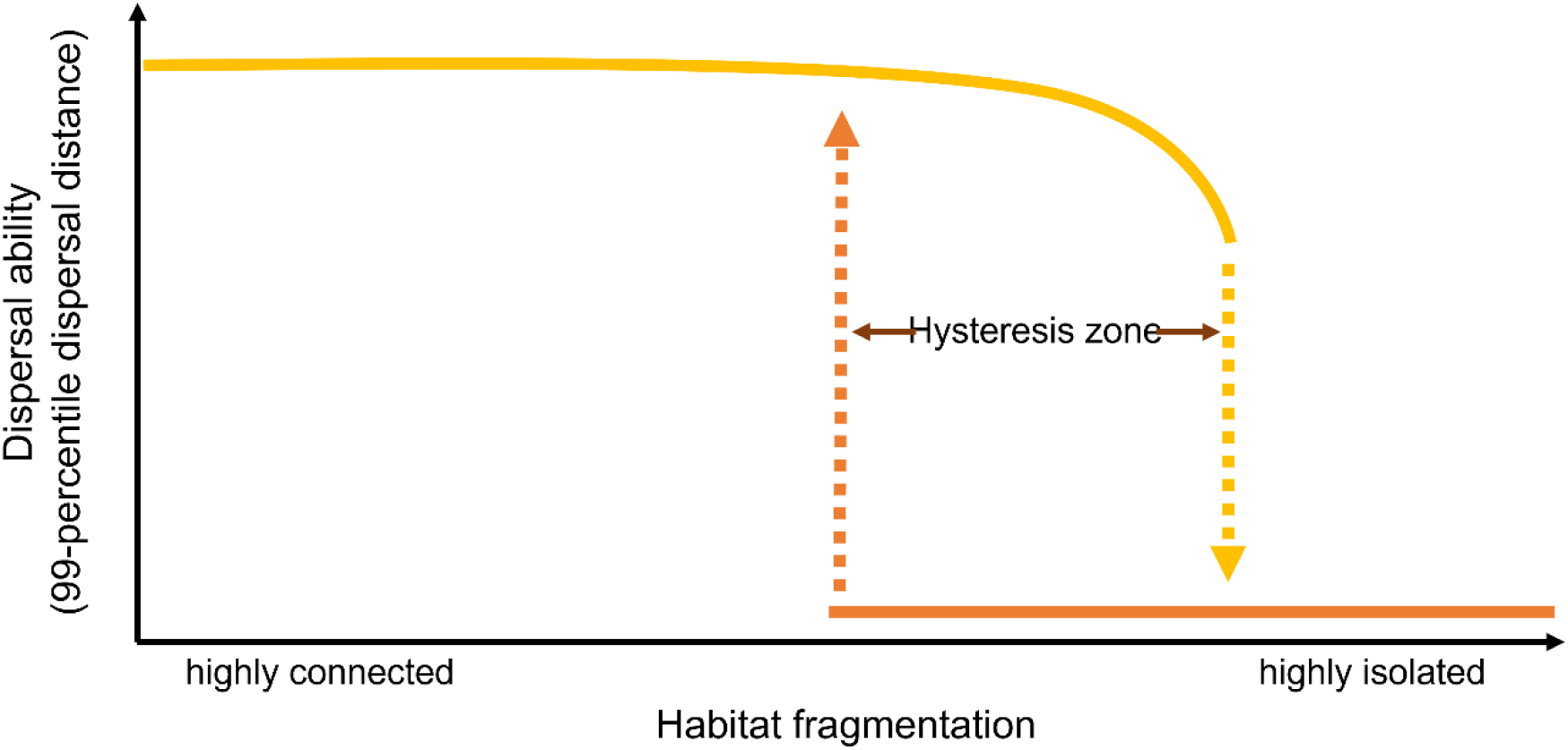
Hypothesized effect of habitat fragmentation on dispersal strategy, taking plant seed dispersal via water as an example. The yellow line indicates evolution of dispersal ability with increasing levels of habitat fragmentation; the orange line shows the evolution of dispersal ability with decreasing habitat fragmentation levels. In the hysteresis zone, the dispersal ability that evolves depends on the previous state of the system.

Collecting empirical evidence establishing the hypothesized evolutionary trajectories would involve experimental habitat manipulation over many generations, which would require an extensive period of time that is not available given the urgency of mitigating effects of habitat fragmentation. Therefore, we use a simple model, based on a well-documented and data-rich system: dispersal by water of shoreline plant species. Dispersal of shoreline plant seeds by stream or river water flow provides a well-studied, mostly unidirectional and linear study system: shoreline plants produce seeds that are released onto, or taken up by, stream or river water, which transports them downstream until they are deposited on the bank (Soons et al. 2017; de Jager et al. 2019; Nilsson et al. 2010; Merritt and Wohl 2002). Recent advances established (i) that shoreline plants have long-floating seeds adapted to dispersal by surface water flows as primary dispersal mechanism, securing eventual deposition at the shoreline (Soons et al. 2017), and (ii) that larger seeds are dispersed over longer distances by river water flow, following a simple exponential relation between seed size and dispersal distance (de Jager et al. 2019), while (iii) upstream dispersal occurs via other, usually accidental, vectors to which plant species do not have clear adaptations (Wubs et al. 2016). Riparian plant habitat is fragmented (Lake 2000) and ongoing habitat degradation and restoration both further affect this heterogeneity (Lake et al. 2007). We used this simple system to provide a theoretical proof-of-principle of our hypothesized hysteresis effect in dispersal evolution as a consequence of habitat fragmentation and restoration events.

## Methods

To simulate dispersal of shoreline plant seeds by water and assess the potential for evolution of dispersal strategies in fragmented landscapes, we created a simple, computationally fast, deterministic model, and a more realistic, stochastic, individual-based model. In both simulation models, we used seed size as the mutable trait under selection and the colonized habitat area (expressed in average number of grid cells colonized by a plant’s seeds) as the measure of fitness to be maximized. In the deterministic model, we simulated seed dispersal by one individual in an otherwise empty environment, whereas in the stochastic model, we simulated seed dispersal by multiple individuals.

We simulated semelparous seed dispersal in a unidirectional, linear system consisting of habitat patches, consisting of 50 grid cells per patch (*X*_*H*_ = 50), separated by inter-patch distances (*IPDs*) that contain non-habitable patches, consisting of *X*_*IPD*_ cells per patch. A grid cell can contain only one plant individual. Between habitat patches, we first increased inter-patch distances (*X*_*IPD*_) from 0 to *X*_*max*_ grid cells between model runs, and subsequently decreased the inter-patch distance again from *X*_*max*_ grid cells back to 0, while using the seed sizes evolved in the previous run as the initial seed sizes in the next run to examine hysteresis effects.

Seed size was defined by seed length, which varies for shoreline plants (helophytes) between 0.5 and 48 mm, with 95% of the seed lengths between 0.5 and 27 mm (Kleyer et al. 2008; de Jager et al. 2019). The number of seeds dispersed depends on the seed size, as larger seeds are costlier to produce (see below).

Seed dispersal by water depends on the probability that a seed becomes entrapped on a river or stream bank. Assuming no differences in entrapment probability between habitat patches and uninhabitable areas, the fraction of seeds remaining in the water column after dispersing a distance *d* follows an exponential decay function:

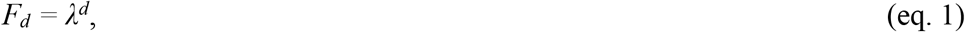

where *λ* is the decay exponent (de Jager et al. 2019). Following de Jager *et al*. (2019), we assume that *λ* is a sigmoidal function of seed size (*S*):

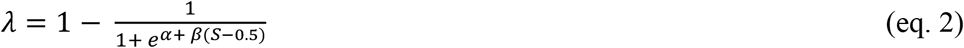

where *α* and *β* define the slope of the sigmoid curve (*β*) and the point where *λ* = 0.5 (equal to *α/β*). The parameters *α* and *β* can be calculated from the median dispersal distances (*d*_*0*.*5*_ and *d*_*30*_ at *F*_*d*_ = 0.5) of the smallest and largest seed sizes considered in our model (0.5 and 30mm, respectively; Suppl. Methods A):

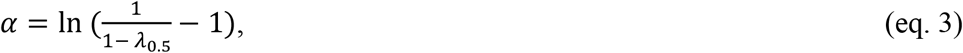

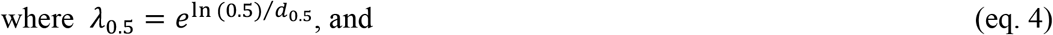

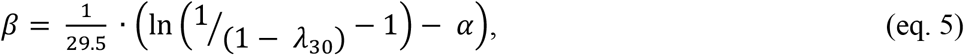

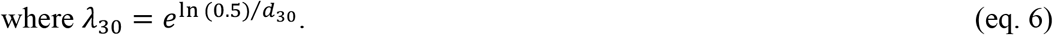

### Deterministic model

Per habitat cell, we calculated the probability that at least one of the seeds dispersed to this cell germinates and survives. Combining the colonization successes at all habitat cells, the summed probabilities of all habitat cells becoming occupied by a plant’s seeds determines the fitness of the simulated plant. For a given seed size *S*, the number of seeds dispersed to a grid cell at distance *x* from the parental patch is given by:

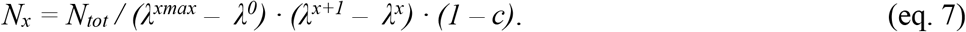

Here, *N*_*tot*_ is the total number of seeds to disperse, *xmax* and 0 are the edges of the simulated environment (*xmax* = 50,000), and *c* is the production cost parameter:

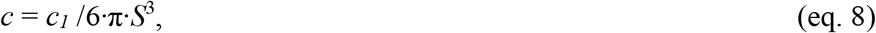

where the cost was calculated per seed volume with cost parameter *c*_*1*_ = 0.0001. We assume (i) round seeds, (ii) that large seeds are more costly to produce, and (iii) that large seeds have a higher chance of survival. This trade-off between production cost and survival chance is incorporated in *c* to simplify the model. Naturally, the shape of the seed size cost function will affect simulation results. In our sensitivity analysis, we vary the value of *c*_*1*_ to examine its effect.

In the deterministic model, we did not include any form of competition, as only one plant individual was simulated per run. The only competition that was implicitly included was between kin: of all the seeds that a plant dispersed to the same grid cell, only one could survive. Given a certain probability of germination and survival, *g*, the probability that a grid cell will be occupied (*P*^*+*^) depends on the number of seeds dispersed to it:

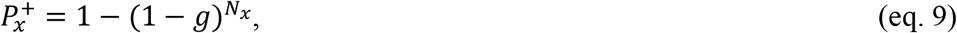

where *N*_*x*_ is the number of seeds dispersed to the grid cell at distance *x* from the parental grid cell. In the current model, we set *g* to 0.3 for habitat patches and 0 for uninhabitable patches. In our sensitivity analysis, we varied the value of *g* to examine its effects. We used the sum of 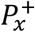 between 0 ≤ *x* ≤ *xmax* grid cells as a measure of fitness, as it indicates the average number of grid cells and thus total habitat area that may be colonized by a plant’s seeds. We compared these fitness measures between plants that produce seeds of a certain size *S* (0.5 < *S* ≤ 30 mm) to fitness of plants that produce slightly larger or smaller seeds (*S*^*-*^*= S – 0*.*01 mm* and *S*^*+*^ *= S +*0.01 mm). When fitness of S^-^ or S^+^ is higher than that of S, seed size evolved and new comparisons were made between the new fitness values, until an equilibrium *S*^*\**^ was reached. With this model, we thus disregard the time it takes for an equilibrium seed size to evolve, the probability that multiple seed sizes may evolve simultaneously, and stochasticity in fitness.

With the seed size evolved in an environment with inter-patch distance *X*_*i*_ (starting with *X*_*0*_ = 0), we initiated seed size evolution in the environment with inter-patch distance *X*_*i+1*_, until *X-* _*max*_ (*X*_*max*_ = 5000) was reached, after which we decreased inter-patch distance again to 0, one grid cell at a time. All simulations were run once; since our model is deterministic, there is no need for replications of simulation runs.

### Stochastic model

In the stochastic model, boundaries are continuous. Initially, simulations start with *X*_*IPD*_ = 5m and all habitable cells hold a plant that produces seeds with intermediate seed size *S* = 20mm. Plants one by one (in a random order) disperse their seeds, until either all seeds are dispersed or all habitable cells are occupied by the next generation of seedlings. The number of seeds dispersed by plant *j* depends on seed size and the parameter *N*_*tot*_:

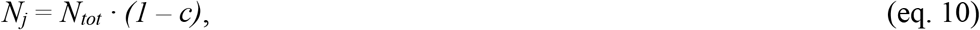

where *c* is calculated as in eq. 8. Dispersal distance of the randomly selected seed is drawn from an exponential distribution (eq. 1). If the seed disperses to a habitable cell that is unoccupied by a seedling (ignoring the parent generation), it can germinate and survive there with probability *g*. Seeds dispersed to inter-patch locations or to cells already occupied by other seedlings have zero survival and germination probability. Established seedlings can have a larger or smaller seed size (+ or – 1mm) than their parent with a mutation rate *μ* = 0.001. After a fixed number of generations (*N*_*gens*_), we recorded the population’s seed size distribution and increased the inter-patch distance with 5m (i.e. 5 cells). When *X*_*IPD*_ = 1,000m was reached, we again decreased the inter-patch distance with 5m per *N*_*gens*_ generations. This model thus simulates evolution of dispersal distances and includes stochasticity and competition, but is computationally much more demanding, which is why it was run for a more limited set of conditions.

### Sensitivity analysis

As model results may depend on parameter values, we ran additional simulations varying the parameters patch size (10, 50, 100, and 500 grid cells), survival and gemination (*g* = 0.9, 0.6, 0.1, and 0.01), seed size cost (*c*_*i*_ = 0.000001, 0.00001, 0.001, and 0.01), total number of seeds to disperse (*N*_*tot*_ = 100, 1000, and 100,000), number of generations per inter-patch distance (*N*_*gens*_ = 10, 100, 1000, and 10,000), and mutation rate (*μ* = 0.01, 0.001, 0.0001).

## Results

Results from our deterministic and stochastic model confirm that, at least in theory, dispersal capacity can be abruptly lost once habitat patches become too far apart and, likewise, can be gained abruptly again once the distance between such patches is sufficiently decreased (Fig. 2, 3). In these cases, there is a hysteresis zone (Fig. 2, 3). However, such responses are not the case, for the entire parameter space (Fig. 4; Fig. 5): for some parameter combinations, there is a gradual decrease of mean dispersal distance with increasing distances between habitat patches, and a gradual decrease back again with decreasing inter-patch distances. For the deterministic model, the size of the hysteresis effect increased when the difference in dispersal distances between small and large seeds increased (i.e., when larger seeds dispersed much farther than smaller seeds, the hysteresis effect was strongest). For the stochastic model, the size of the hysteresis effect was mostly large when both small and large seeds dispersed over long distances (i.e., when larger seeds dispersed over long distances relative to the patch size, and small seeds also dispersed over reasonable distances, the hysteresis effect was strongest). In most simulations, especially those with the stochastic model, dispersal capacity first increases with inter-patch distances, before it is abruptly lost (Fig. 2, 3).

**Figure 2:**
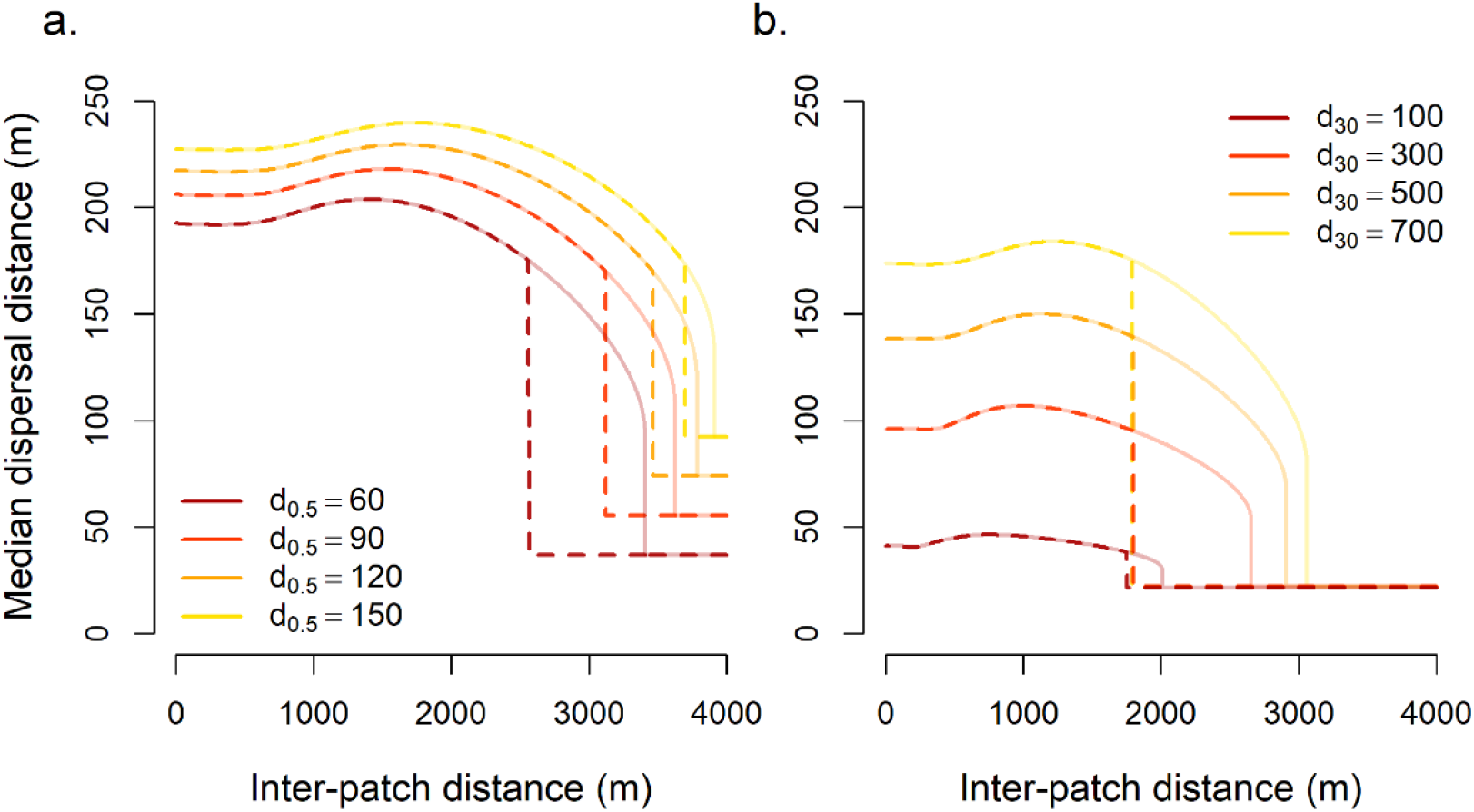
Examples of evolved median dispersal distances in simulations with the deterministic model with increasing (solid lines) or decreasing (dashed lines) inter-patch distances (IPD). In (a), d_30_ = 700; in (b), d_0.5_ = 30. Other parameter values were kept constant at X_H_ = 50, N_tot_ = 10,000, g = 0.3, and c_1_ = 0.0001.

**Figure 3:**
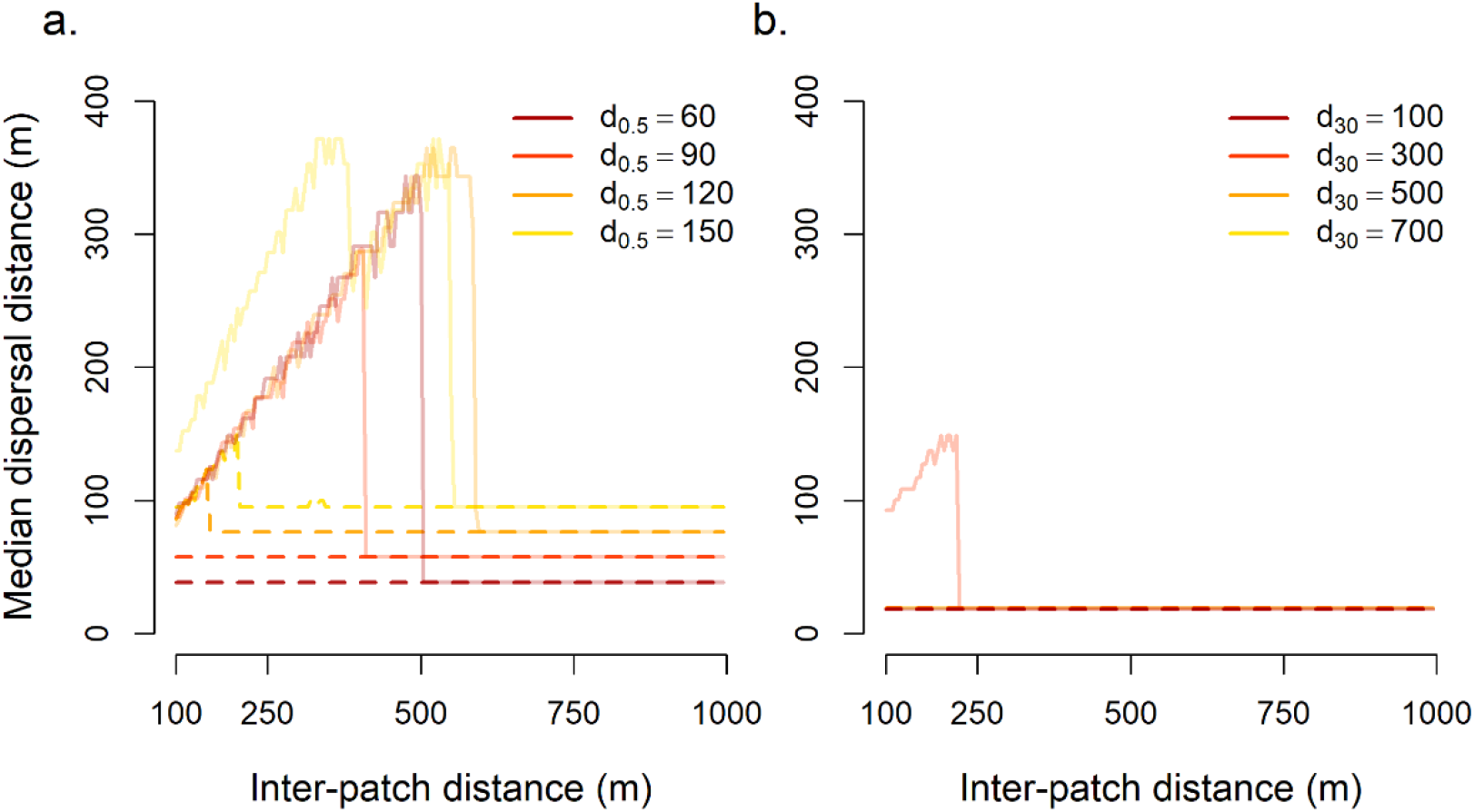
Examples of evolved median dispersal distances in simulations with the stochastic model with increasing (solid lines) or decreasing (dashed lines) inter-patch distances (IPD). In (a), d_30_ = 700; in (b), d_0.5_ = 30. Other parameter values were kept constant at X_H_ = 50, N_tot_ = 10,000, g = 0.3, c_1_ = 0.0001, N_gens_ = 10,000, and μ = 0.001.

**Figure 4:**
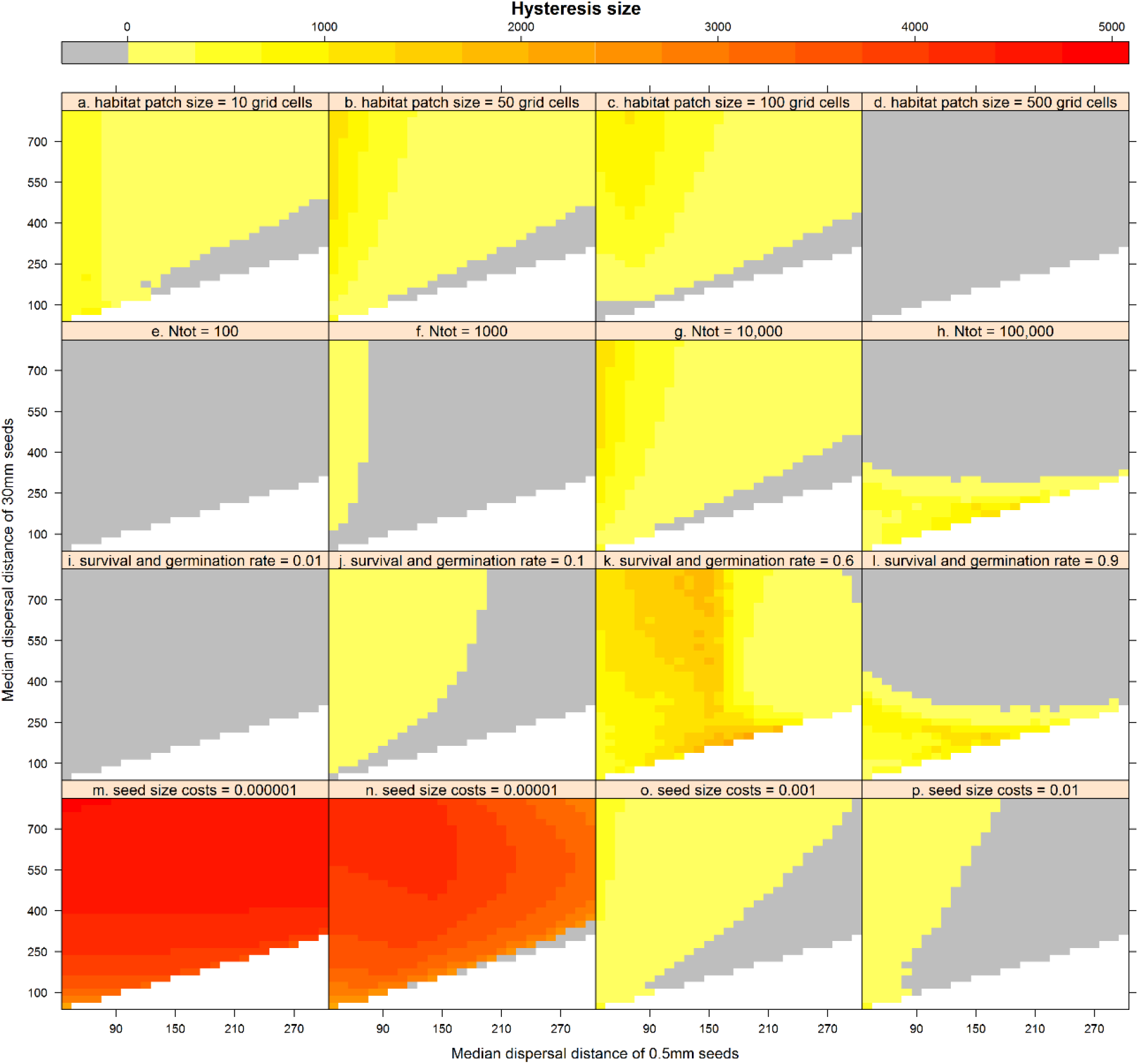
Size of the hysteresis area (in cells), using the deterministic model, for a range of parameter value combinations, varying the median dispersal distance of 0.5mm seeds (in cells), the median dispersal distance of 30mm seeds (in cells), and the habitat patch size (a-d), the total number of seeds (e-h), survival and germination rate (i-l) or the seed size costs (m-o). Grey areas indicate no hysteresis zone, white areas contain no data (d_0.5_ ≤ d_30_). If not the changing parameter, parameter values used in simulations are X_H_ = 50, N_tot_ = 10,000, g = 0.3, and c_1_ = 0.0001.

**Figure 5:**
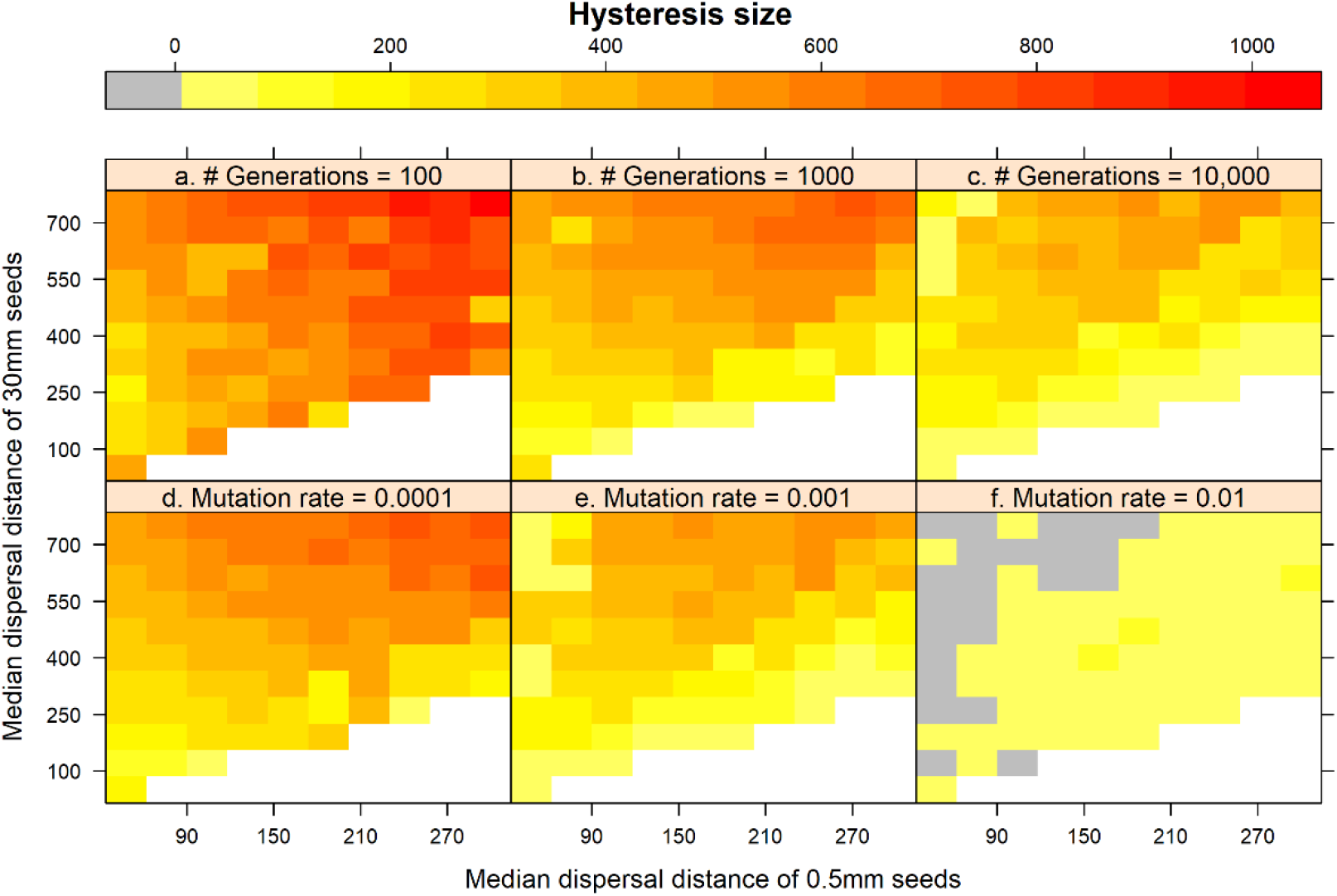
Size of the hysteresis area (in cells), using the stochastic model, for a range of parameter value combinations, varying the median dispersal distance of 0.5mm seeds (in cells), the median dispersal distance of 30mm seeds (in cells), and the number of generations per level of habitat fragmentation (a-c) or the mutation rate (d-f). Grey areas indicate no hysteresis zone, white areas contain no data (d_0.5_ ≤ d_30_). If not the changing parameter, parameter values used in simulations are X_H_ = 50, N_tot_ = 10,000, g = 0.3, c_1_ = 0.0001, N_gens_ = 10,000, and μ = 0.001.

Our sensitivity analyses indicate that, for the deterministic model, no hysteresis was observed when habitat patch sizes were large (*X*_*H*_ = 500 cells), the total number of seeds small (*N*_*tot*_ < 1000), or the survival and germination rate small (*g* = 0.01). When habitat patches are large, long-distance dispersal will remain a good strategy, as little seeds are lost to the uninhabitable matrix between patches. Conversely, short-distance dispersal is the best strategy when the total number of seeds dispersed is small or the germination rate low, as there is little kin competition in these cases. The largest hysteresis areas were observed when seed size costs were small (*c*_*1*_ ≤ 0.00001). Here, seed size and thus dispersal capacity is maximized with increasing inter-patch distances, until a critical threshold is reached, after which the smallest possible seed size – with the shortest dispersal distances – is most efficient, as the least seeds are lost to the uninhabitable matrix between patches. Simulations with the stochastic model resulted in larger hysteresis areas when time between changes in inter-patch distances were short (*N*_*gens*_ = 100) or mutation rates small (*μ* = 0.001), e.g. when there is insufficient time between environmental changes to adapt to the new situation.

## Discussion

Our model suggests that, for species in a very simple system consisting of habitable patches separated by a non-inhabitable matrix, whose movements are non-directed and across distances primarily determined by organism traits, increasing habitat fragmentation of relatively small habitat patches may result in an abrupt loss of dispersal ability that is extremely difficult to reverse. When the distances between habitat patches are small, seed dispersal over long distances evolves. This is in line with general theoretical predictions of dispersal, where long-distance dispersal strategies are favored in continuous habitats or habitable areas separated only by short distances, due to the avoidance of kin competition (Treep et al., 2021). With increasing distances between habitat areas, seed dispersal capacity first increases and then decreases gradually. By increasing dispersal capacity, subsequent habitat patches may still be reached when inter-patch distances increase. Yet, when distances between patches become too large, seed dispersal capacity decreases. From this point on, loss of seeds dispersing into the unsuitable matrix is counterbalanced by dispersing more seeds within the parental area. Then, at a certain critical distance between habitable areas, even the slightest additional increase in this distance results in a collapse in seed dispersal capacity. Interestingly, a similar non-linear relation between dispersal capacity and landscape fragmentation has also previously been shown in an earlier theoretical study, which demonstrated that connectivity may be abruptly lost if model organisms gradually decrease their dispersal capacity (Keitt et al. 1997).

A crucial result from our simulations is that the distance between habitat patches may need to be greatly reduced to effectively restore movement between habitat fragments. In other words, the connectivity between patches needs to be restored to a relatively high level in order to allow an evolutionary trajectory towards frequent long-distance dispersal events. As mutations result in small changes in a trait such as seed size, evolution of seed size may not occur through fitness valleys, resulting in a hysteresis zone in seed size. When the distance between habitat patches is decreased but the system remains within the hysteresis zone, short-distance dispersal strategies will remain the evolutionary attractor, as small increases in seed size, and thereby in dispersal distance, result in lower relative fitness (Suppl. Fig. 1). Once the distance between habitat patches has been effectively reduced, the valley in the fitness landscape is no longer present, and small increases in dispersal capacity result in increases in fitness. This process leads to selection of seeds that can disperse over longer distances, effectively reconnecting habitat across the landscape.

Hysteresis in dispersal evolution may explain lack of restoration success due to failure of species to recolonize restored areas in heavily fragmented landscapes where remnant source populations still survive but have been isolated for a relatively long period of time. As biodiversity is under threat throughout the world and populations are being increasingly fragmented (IPBES 2019), it is of critical importance to restore connectivity between habitat patches so that species can overcome possible hysteresis effects and disperse again between habitat fragments. Our study shows that these connectivity levels may need to be much higher than minimal connectivity levels maintaining dispersal in reference populations in relatively pristine landscapes.

Critical transitions from long-to short-distance dispersal in natural populations have not yet been observed, mainly due to the suddenness of the shift and the absence of empirical and experimental studies investigating the effect of increasing habitat fragmentation on dispersal evolution over time. For plant species, Cheptou *et al*. (2008) detected a substantial difference in dispersal capacity between *Crepis sancta* populations in a city and a less fragmented rural area, where short-range dispersal seemed to have evolved in the more fragmented landscape of the city (Cheptou et al. 2008). This study also illustrated one of the mechanisms by which rapid loss of dispersal may evolve: a shift in the ratio between two distinct types of propagules, one with appendages to enhance dispersal and one without, on the same plant. Many plant species (>50 genera) within the Asteraceae family exhibit such a seed dimorphism (Imbert 2002), suggesting that rapid evolution of dispersal may be common in populations of this large group of generally wind-dispersed plant species and that species from this family might be a suitable model system to record the first experimental evidence for critical transitions in dispersal.

We selected a well-studied model system that is characterized by unidirectional dispersal with an exponential dispersal kernel. The simplified setup that we chose for our first theoretical proof-of-principle of evolutionary hysteresis in dispersal capacity can be improved and extended upon in future studies. For instance, we created a one-dimensional environment, while most dispersal occurs in two-dimensional habitats. Expanding the model used in Treep *et al*. (2021) may show that the critical transitions we observe in our simulated 1-D environments can also occur in 2-D environments. Also, while our simulated habitat patches were of equal quality throughout the modelled space, habitat quality is highly diverse in real landscapes. In fact, reduction of habitat quality rather than a complete loss of habitat is generally underlying habitat degradation and fragmentation. In future studies, one might include variation in habitat quality when modelling eco-evolutionary dynamics of seed dispersal (Mortelliti et al. 2010; Liao et al. 2013*b*). Also, we placed habitat patches equidistantly from another. Yet, clustering of habitat patches is highly likely to result in different ecological and evolutionary outcomes (Hiebeler 2000; Liao, Li, Hiebeler, Iwasa, et al. 2013). In such heterogeneous environments, multiple dispersal strategies can coexist, and community composition can vary substantially between locations (Hiebeler 2007; Liao, Li, Hiebeler, El-Bana, et al. 2013; Liao, Li, Hiebeler, Iwasa, et al. 2013; Cote et al. 2017). An extension of our model may show us the effects of spatial heterogeneity on dispersal evolution.

Our model is based on an exponential dispersal kernel, which is typical for seeds dispersed by water. Many plant species, across a range of dispersal mechanisms, have even more leptokurtic dispersal distributions that are characterized by fat tails (Bullock et al. 2017). However, in plant species that are dispersed by vertebrates, the navigation capacity of the vector may play an important role in seed dispersal, especially in fragmented landscapes where large vertebrates are able to maintain connections between otherwise isolated patches (Kleyheeg et al. 2017). Such species are likely to show a very slow evolutionary trait trajectory towards limited or restored movement and may lack a hysteresis in response to defragmentation. Also, including habitat dynamics (i.e., turnover of habitable areas) is likely to significantly affect the outcomes of our model. Slow habitat dynamics (i.e., patch life span much exceeding organism life span) will result in evolution of short-distance and long-distance dispersal in much the same way as in permanent habitable areas (Treep et al., 2021). However, long-distance dispersal is essential for survival in highly dynamic habitats (Treep et al., 2021). If habitat fragmentation in such environments exceeds a certain threshold, evidence suggests that population extinction rather than loss of dispersal is the likely outcome (Treep et al., 2021; Fountain et al. 2016). An empirical study on extinct Glanville fritillary butterfly populations shows that dispersal capacity was enhanced instead of inhibited in response to increased habitat fragmentation, before regional collapse of dispersal capacity and extinction of the species in the most disturbed regions occurred (Fountain et al. 2016). Some of our results of the stochastic model show the same patterns as observed in this study. In this species, the sudden collapse of dispersal capacity and extinction of the population coincided, which is in line with the result of model simulations with high habitat dynamics (Treep et al., 2021). Based on the exploratory work presented here, we suggest that new lines of research should include unravelling the relations between habitat fragmentation and dispersal limitation in (i) organisms with directed dispersal and (ii) in highly dynamic habitats, besides the (iii) empirical testing of our hypotheses in existing and experimental populations. It is of urgent importance to extend this research and find further validation and empirical support to prove that our proposed mechanism is responsible for the observed discrepancies between habitat restoration and actual colonization, thereby providing new conservation strategies that are needed to overcome the ecological and evolutionary consequences of habitat fragmentation.

## Author contributions

MdJ conceived the hypotheses, created the model, and analyzed its results. MdJ and MS discussed the results and implications and wrote the manuscript.

## Data availability

The datasets generated and analysed during the current study are available from the corresponding author on request.

## Code availability

A C++ script containing the function to calculate the number of grid cells with germinated and surviving seedlings is available as a Supplementary File.

