## Supplementary Methods and Figure for "Critical transitions and evolutionary hysteresis in movement: Habitat fragmentation can cause abrupt shifts in dispersal that are difficult to revert"

By M. de Jager & M.B. Soons

Contents:

Supplementary Methods A p2

Supplementary Figure 1 p3

**Supplementary Methods A**

The parameters $\alpha$ and $\beta$ can be calculated from the median dispersal distances ($d_{0.5}$ and $d_{30}$) of the smallest and largest seed sizes considered in our model (0.5 and 30mm, respectively). When seed size $S$ = 0.5mm, eq. 2 results in

$$\lambda = 1 - \frac{1}{1+e^{\alpha+\beta(0.5-0.5)}} = 1 - \frac{1}{1+e^{\alpha}} . \quad \text{(A1)}$$

Reorganizing this equation gives

$$\frac{1}{1+e^{\alpha}} = 1 - \lambda, \quad \text{(A2)}$$

$$1 + e^{\alpha} = \frac{1}{1-\lambda}, \quad \text{(A3)}$$

$$e^{\alpha} = \frac{1}{1-\lambda} - 1, \text{ and} \quad \text{(A4)}$$

$$\alpha = \ln\left(\frac{1}{1-\lambda} - 1\right). \quad \text{(A5)}$$

As the median dispersal distance occurs at $F_d$ = 0.5, we can rewrite eq. 1 to eq. 3. When seed size S = 30mm, eq. 2 results in

$$\lambda = 1 - \frac{1}{1+e^{\alpha+\beta(30-0.5)}} = 1 - \frac{1}{1+e^{\alpha+29.5\beta}} . \quad \text{(A6)}$$

Reorganizing this equation gives

$$\frac{1}{1+e^{\alpha+29.5\beta}} = 1 - \lambda, \quad \text{(A7)}$$

$$1 + e^{\alpha+29.5\beta} = \frac{1}{1-\lambda}, \quad \text{(A8)}$$

$$e^{\alpha+29.5\beta} = \frac{1}{1-\lambda} - 1, \quad \text{(A9)}$$

$$\alpha + 29.5\beta = \ln\left(\frac{1}{1-\lambda} - 1\right), \quad \text{(A10)}$$

$$29.5\beta = \ln\left(\frac{1}{1-\lambda} - 1\right) - \alpha, \quad \text{(A11)}$$

$$\beta = \frac{1}{29.5} \cdot \left(\ln\left({}^{1}/_{(1-\lambda_{30})} - 1\right) - \alpha\right), \quad \text{(A12)}$$

As the median dispersal distance occurs at $F_d$ = 0.5, we can rewrite eq. 1 to eq. 6.

**Supplementary Figure 1**

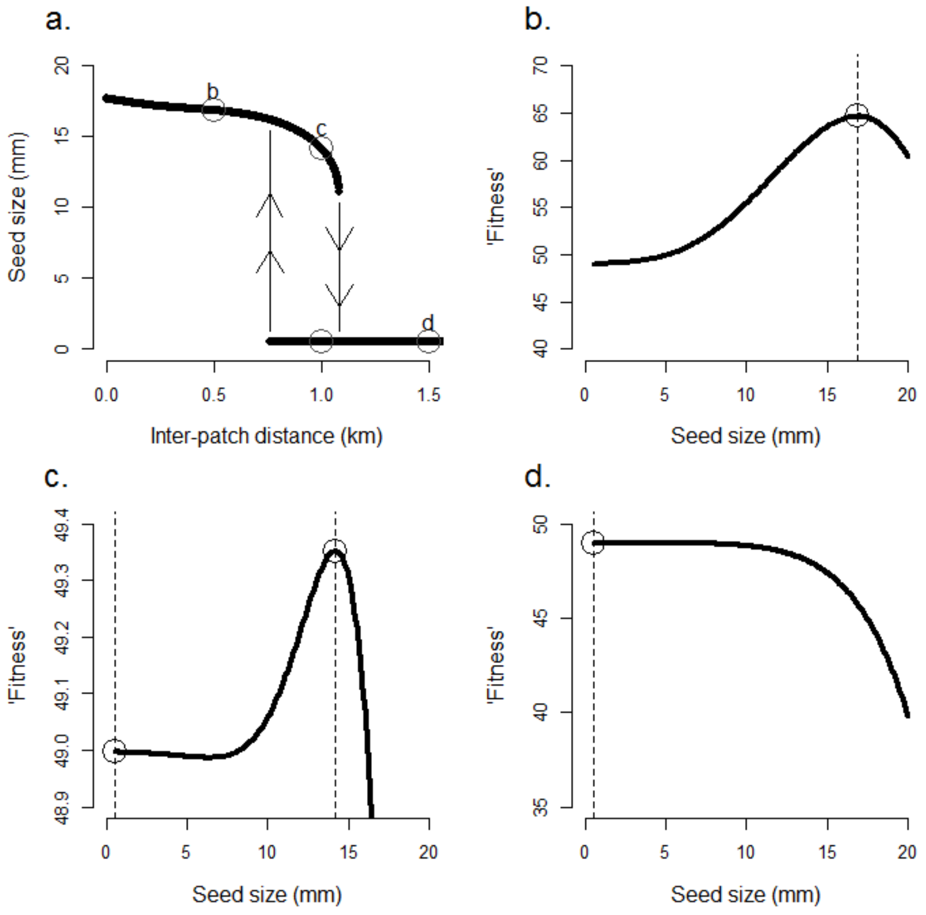


***Supplementary Figure 1:*** *Evolution of evolutionarily stable dispersal strategy in relation to habitat fragmentation, with evolutionary stable strategies per distance between patches in (**a**). Optimal seed sizes based on number of occupied grid cells are illustrated for three different fragmentation levels, represented by distances between habitat patches of 500 (**b**), 1,000 (**c**), and 1,500 (**d**) grid cells (0.5, 1.0, and 1.5 km, respectively). Open circles in (**a**) indicate how these are translated to the hysteresis graph, where two different dispersal strategies can occur when a fitness valley is present in the relation between number of occupied grid cells and seed size (**c**). Parameter values used in these simulations are* $X_H = 50$, $N_{tot} = 10{,}000$, $g = 0.1$, $c_1 = 0.0001$, $d_{0.5} = 40$ *and* $d_{30} = 750$.
